# Assessment of the household antibiotic resistance genes, virulence factor genes, and pathogen profiles from three global cities

**DOI:** 10.64898/2026.04.09.717483

**Authors:** Caroline Scranton, Victoria Obergh, Madison Goforth, Kavitha Ravi, Poornima Jayakrishna, Anisa S.K., Karl Krupp, Purnima Madhivanan, Stephanie Boone, Charles Gerba, Frank Xu, M. Khalid Ijaz, Kerry K. Cooper

## Abstract

This study assessed the prevalence of antibiotic resistance genes (ARGs) and virulence factor genes (VFs), DNA viruses, and medically-relevant pathogens in three major cities around the globe – Mysuru (India), Dubai (United Arab Emirates), and Tucson (Arizona, United States of America). Ten households were sampled in each city, at ten sites in the bathroom, kitchen, and living spaces. The alpha diversity of ARGs significantly differed between household locations in each country (ANOVA, p<0.05) and beta diversity (dissimilarity) analysis showed a significant association between the ARGs and the geographic locations (PERMANOVA, p<0.01). A set of ARGs were found in every location across the dataset (the core ARG profile) included 25 different genes. The alpha diversity of virulence factors differed across the household locations within the three cities (ANOVA, p<0.01). The beta diversity of VFs was not well explained by geographic location or location within the household (PERMANOVA, p=0.129 (geographic), p=0.127 (household)). There were 341 unique VFs found in the study, but only 5 core VFs across the dataset. Bacterial pathogens detected across the household included *Escherichia coli*, *Acinetobacter baumanii*, *Klebsiella pneumoniae*, and more. The DNA (and bacteriophage) virome varied between countries and was more diverse in Tucson and Dubai (top viral families included Poxviridae and Orthoherpesviridae - two families which contain human pathogens - and Steitzviridae, a family of bacteriophages) compared to Mysuru, where nearly all viruses were a part of the Muvirus genus (a bacteriophage which contributes to horizontal gene transfer by phage transduction).

**Importance:** The diversity of the built environment microbiome is not well characterized globally. Household occupants interact with the built microbiome on a daily basis, and the built microbiome may have the potential to influence human health. The presence of pathogens in the built environment and the key genes contributing to microorganism pathogenicity were investigated in this study, as information on this is lacking on an international scale. The diversity of ARG and VFs across the globe, as well as the presence of medically relevant pathogens within the house that were found in this study highlights the need to explore further the factors which influence the household microbiome, virome, and resistome. This may aid in identify how the build microbiome may be shaped by humans and influence human health.

**Impact Statement:** This research contributes to the understanding of the built microbiome, specifically how it varies within the house, within cities, and across the globe. This can aid in our understanding of microbial dynamics in environments with heavy human influence and help develop and improve hygiene habits and products which are mindful of the existing microbiome.

## Introduction

The microbiome of built locations is complex and variable and has the potential to impact human health in a variety of ways(1–3). As a society, we spend increasing time indoors, particularly in areas with higher levels of socioeconomic development and harsh climate, the influence of individuals on the household microbiome (and vice versa) grows(3–7). Previous household microbiome studies have focused on the taxonomic composition across the house; less research has been conducted on the presence of pathogenic microbes and on the genes which are carried within the microbiome, such as antibiotic resistance genes (ARGs) and virulence factors (VFs). Though some species of bacteria are frank pathogens and others opportunistic, many bacteria maintain the ability to uptake and utilize ARGs or VFs to drive their survival by horizontal gene transfer and/or mobile genetic elements (MGEs) and enhance virulence or disseminate antibiotic resitance(2, 8–10). The diversity and prevalence of such genes is of interest, and understanding the role of these genes is essential to assess the potential impacts of the built microbiome on human health.

Several studies have detected ARGs and VFs in the built environment. Analysis of resistance genes of indoor dust in Beijing, China revealed greater abundances of ARGs than outdoor dust samples, which contained genes for multidrug, bacitracin, and polymyxin resistance on mobile genetic elements(11). The study also found the most common VFs in the dust were related to motility, metabolism, and adhesion and were widespread among the microbiome(11). A separate study of dust samples in Beijing identified several ARGs that conferred resistance to the clinically-important antibiotics vancomycin and macrolide-lincosamide-streptogramin B (MSL-B), which were influenced by the composition of the microbiome(12). In New Delhi, India, household surface samples found multidrug resistance genes were predominant in all samples, followed by aminoglycosides and bacitracin(13). Various VFs were also found in different bacterial species(13). VFs and ARGs detected in household dust have also been significantly associated with the presence of endocrine-disrupting chemicals, which are widely used in personal care products, processed foods, and cosmetics(14). Marshall et al (2012) found that bathroom and kitchen sites which were treated with antibiotic cleaning products retained high titers of bacteria, and that gentamicin resistance was significantly higher in places treated with antibiotic cleaners than without(15). They also found that culturable microorganisms from household surfaces were resistant to antibiotics at rates as high as 45%, with many microorganisms being multi-drug resistant(15). The microbiome, resistome, and virome of unrestricted buildings (such as public buildings and houses) was found to be distinctly different from that of controlled-access buildings (like intensive care units or factory clean rooms)(16). Many previous studies focus on the microbiome and ARGs in dust in the built environment; this study aims to determine the prevalence of these genes on different surfaces around the house, where individuals may have more contact with the microbes (compared to those in dust) through their everyday activities like cooking and using the bathroom. Additionally, these surfaces may be cleaned more frequently with sanitizers as opposed to mechanical cleaning like sweeping, vacuuming, or dusting. We also explored the presence and abundance of virulence genes, which has not been studied as much as the microbiome and ARGs. Characterizing the microbiome alongside the resistome and pathogens of the household is essential to understanding the impacts of the indoor environment on and by the occupants.

The virome of the built environment also has important public health implications. Bacteriophages have been shown to transfer ARGs between bacteria – one study looked at phage across a range of environments and found that 77% of the phage-encoded ARGs they identified were in human associated environments(17). Phage can transfer genes between bacteria through phage transduction, a type of horizontal gene transfer(18). Other types of viruses can also persist in the built environment - indoor air at the University of Colorado in Boulder, CO (USA) was found to harbor DNA and RNA viruses including bacteriophages for Proteobacteria, Firmicutes, and Actinobacteria (some of which are linked to the natural skin microbiome) and viruses (e.g., Papillomaviruses, some members which are responsible for cutaneous and mucosal warts including cervical cancer) which infect eukaryotes, including humans(19). A 2023 study of the built environment virome in Hong Kong noted a habitat (surface type)-dependent distribution of viruses, and that this was the largest driver of viral diversity(20). The Caudoviricetes class was found ubiquitously across the built environment (except for in air samples). It was found to carry a range of Acr proteins, which protect against bacterial CRISPR-Cas9 systems, suggesting a highly coupled host (bacterial) and virus link. This interaction would influence the bacterial microbiome of the built environment(14). The majority of studies focused on viruses in the built environment target the survival of pathogens like SARS-CoV-2, influenza virus, and norovirus(21). Therefore, a knowledge gap exists regarding the virome of the built environment, as it plays a role in the structure of the entire built microbiome and also may have a critical role in the virulence of bacterial pathogens present in the home.

Previous studies have identified potential, opportunistic, and frank pathogens on household surfaces. In Houston, TX (USA), a study of ten households found at least one kitchen surface per household was contaminated with a bacterial foodborne pathogen (*Escherichia coli* was most commonly identified)(22). The previously mentioned study in New Delhi, India also found several pathogenic, opportunistic, or potentially pathogen species that varied across the household(13). The study also identified several DNA viruses in the household virome(13). A study from Boston, MA and Cincinnati, OH (USA) cultured twice as many bacteria and found twice as many species in surface swab samples from kitchens compared to bathrooms(15). Notably, while direct incidence of pathogens were not necessarily identified, studies observing the phylum Actinobacteria are of concern primarily because some species and strains within the phyla exhibit potential for extreme drug resistance(23). Zhou et al found pathogens in indoor dust samples in Beijing were carriers of ARGs and VFs, potentially increasing the risk and severity of human infection by these bacteria(11). The presence of both pathogens and ARGs/VFS in the household environment highlights the potential for severe human infection from the built microbiome – understanding how pathogen and genes are distributed around the house will provide key information on reducing the risk of such infections.

Characterizing the differences in ARGs, VFs, bacterial pathogens, and the DNA virome across a variety of household surfaces and between households of different socio-economic and cultural backgrounds is critical to gaining a better understanding overall of the potential health impacts of the household microbiome. Understanding the communities that comprise the household microbiome, their potential pathogenic properties, and how they interact with the occupants and are influenced by cleaning habits also has implications for public health. Currently, there are only a few household surface microbiome studies in the United States of America (USA), only one recently published in India, and none from the United Arab Emirates (UAE). This study aimed to assess the household microbiome in three geographic locations with populations of at least 500,000 people including Mysuru, (Karnataka, India), Dubai (United Arab Emirates), and Tucson (Arizona, U.S.A), to profile the ARGs and VFs of each geographic and household location and to identify bacterial pathogens and the virome. We hypothesize that there will be significant variation across household surfaces and between the countries in which genes and bacterial and viral taxa are present, potentially driven by surface type or use and overall, by human establishment in the household environment. Multiple sites throughout each household were sampled with the general focus on the kitchen and bathroom. The overall microbiome of household locations both within and between geographic locations was analyzed to determine the prevalence and types of ARGs and VFs, presence of various bacterial pathogens and virome.

## Results

### Global

#### Antibiotic Resistance Genes (ARGs)

We explored the levels of ARGs present in the different household locations among the three cities. Statistically significant differences in the alpha diversity (Shannon index) were found between the samples collected from the ten household locations in each of the three cities individually (ANOVA, p<0.001 for all cities). Homes in Tucson had the highest ARG diversity on the kitchen counters, underneath the toilet rims, showerheads, and kitchen sinks, and lowest ARG diversity in the toilet bowls and on the coffee tables, whereas in Mysuru it was the toilet rims, toilet seats, and under the toilet rims that had the highest ARGs diversity and the showerheads, coffee tables, and kitchen counters had lowest diversity. Dubai had high ARG diversity in the kitchen sinks, toilet bowls, and on the toilet seats, and low diversity on the coffee tables, TV remotes, and kitchen counters (**Figure 1A**). Beta diversity analysis of the ARGs found that the ARG profile of a household microbiome sample is significantly explained by geographic location (PERMANOVA, p=0.002) but not household location (p=0.456) (**Figure 1B**). Core ARGs were determined by identifying ARGs which were found at least once in all houses or in all household locations in each city. Core ARG analysis based on the household location across all three cities found two core ARGs on the coffee tables, eleven core ARGs in both the kitchen and bathroom sinks, eight core ARGs on the toilet seats, nine core ARGs in the toilet bowls, and six core ARGs under the toilet rim. Across the entire dataset, a core set of 22 ARGs against a wide range of antibiotics was identified, meaning at least one household location across the ten houses in each city had one of the twenty-two core ARGs (**Table 1**).

**Figure 1.**
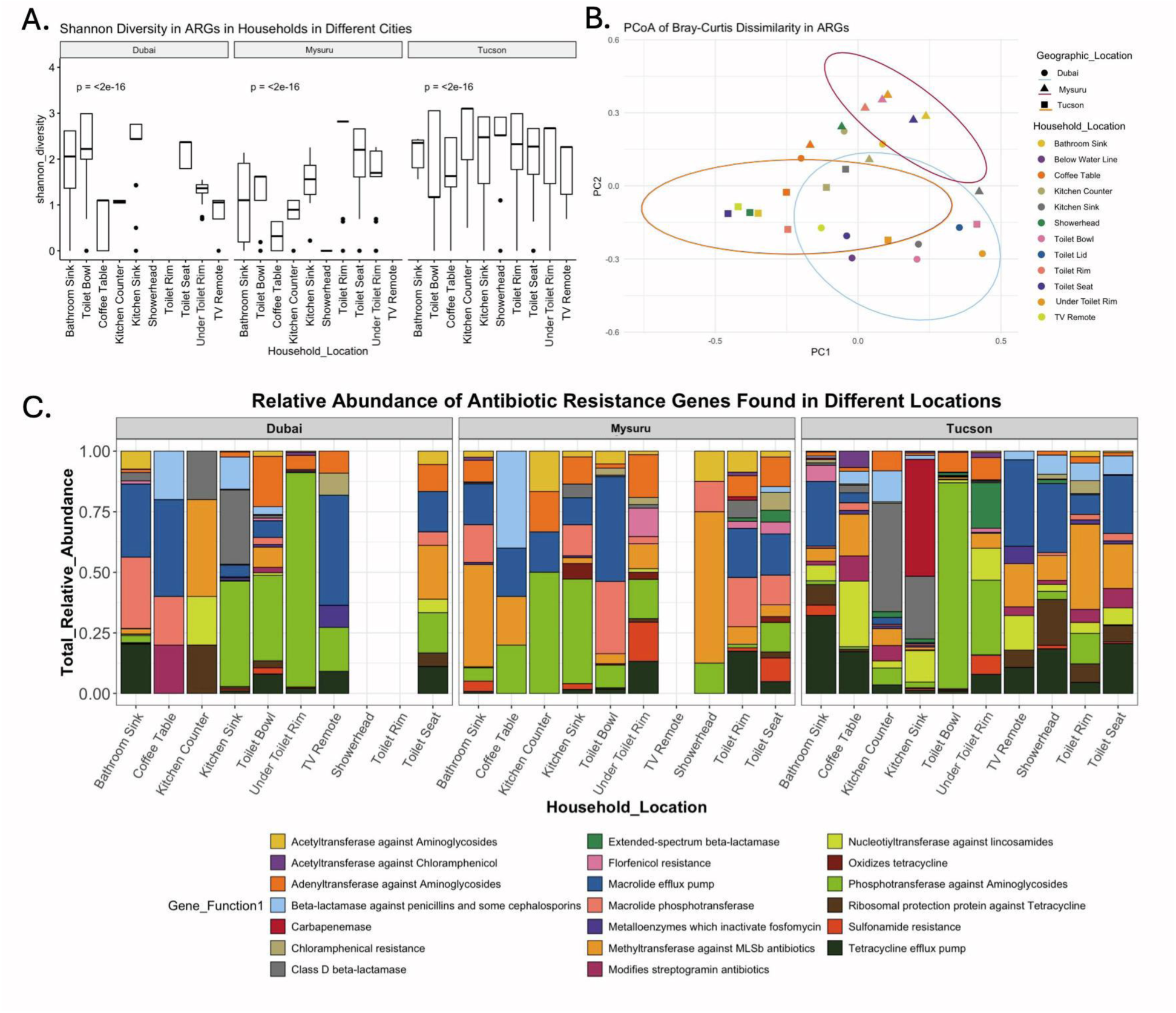
An overview of the antibiotic resistance genes (ARGs) found in the ten household locations across three cities. A: Shannon diversity of ARGs in household locations, significant in Dubai (ANOVA, p<0.05) and highly significant in Tucson and Mysuru (ANOVA, p<0.01). B: Principal coordinates analysis of Bray-Curtis distances (beta diversity) of ARGs in composite household location samples in the study. Samples are colored by location within the house, point shapes correspond to the geographic location, and ellipses are drawn around clusters by geographic location. PERMANOVA analysis showed that ARG diversity is significantly different between geographic locations (p=0.002), but not between household locations (p=0.456). C: The relative abundance of the top 20 most-abundant ARG functions within the composite household samples in the three cities. No ARGs were found in the Dubai composite showerhead or toilet rim samples, or in the Mysuru TV remote sample.

**Table 1.**
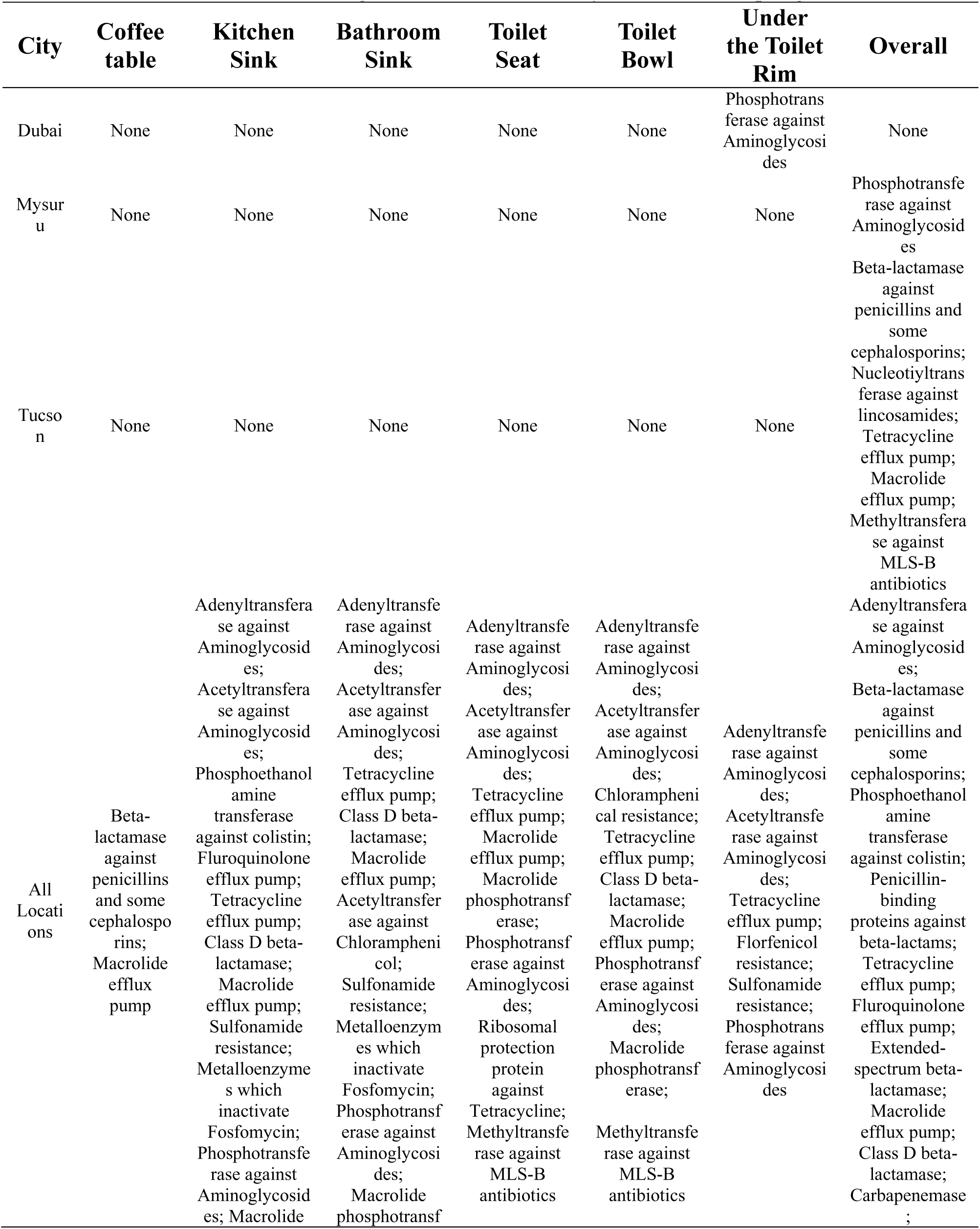
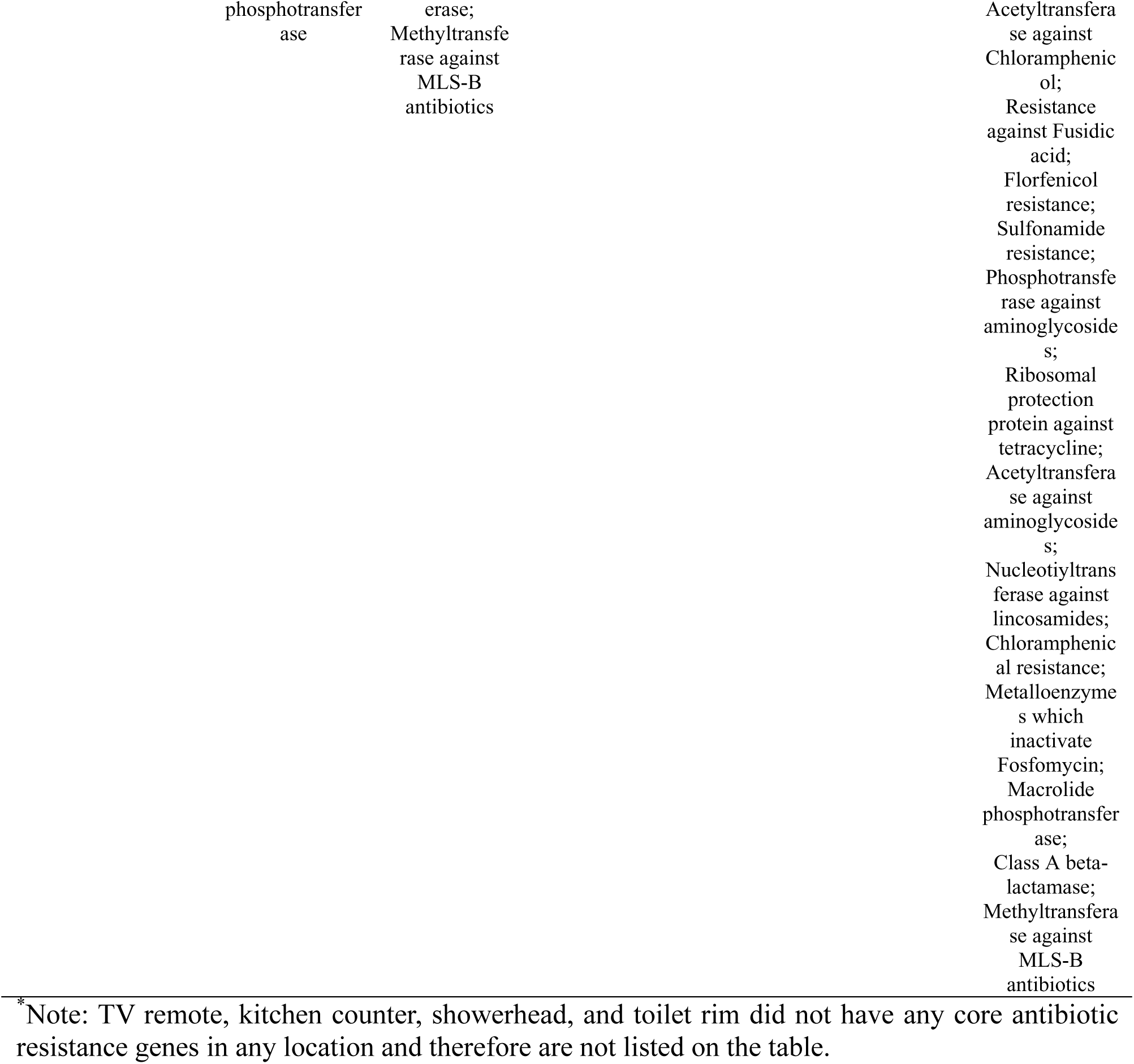
Core antibiotic resistance genes in each location by household sampling site*.

#### Virulence factors (VFs)

We investigated virulence factor (VF) diversity across all the samples and found there was also a highly significant difference (ANOVA, p<0.001) in VF diversity based on household location in all three cities. The locations with the lowest VF diversity were the TV remote, coffee table, and kitchen counter in Dubai and Mysuru, and the coffee table, toilet bowl, and under the toilet rim in Tucson (**Figure 2A**). Based on the Bray-Curtis dissimilarity distance matrix, the beta diversity of VF profiles were not well explained by geographic or household location, with PERMANOVA analysis resulting in non-significant p values (geographic location; p=0.129. household location; p=0.127). Visually, under the toilet rim samples, kitchen sink samples, and coffee table samples clustered more closely together on a PCoA plot compared to the other samples (**Figure 2B**). Core VF analysis was conducted in the same way core ARG analysis was, looking at each household location individually, each house overall, and within each city overall. There were no core VFs in the three cities individually, but across the entire dataset there were five different core VFs identified, meaning that these VFs were found at least once in each city. The core VFs for the individual household locations across the three cities found only flagellar motility genes on all kitchen counters, five core VFs in the kitchen sink, and nine core VFs in the bathroom sink (**Table 2**).

**Figure 2.**
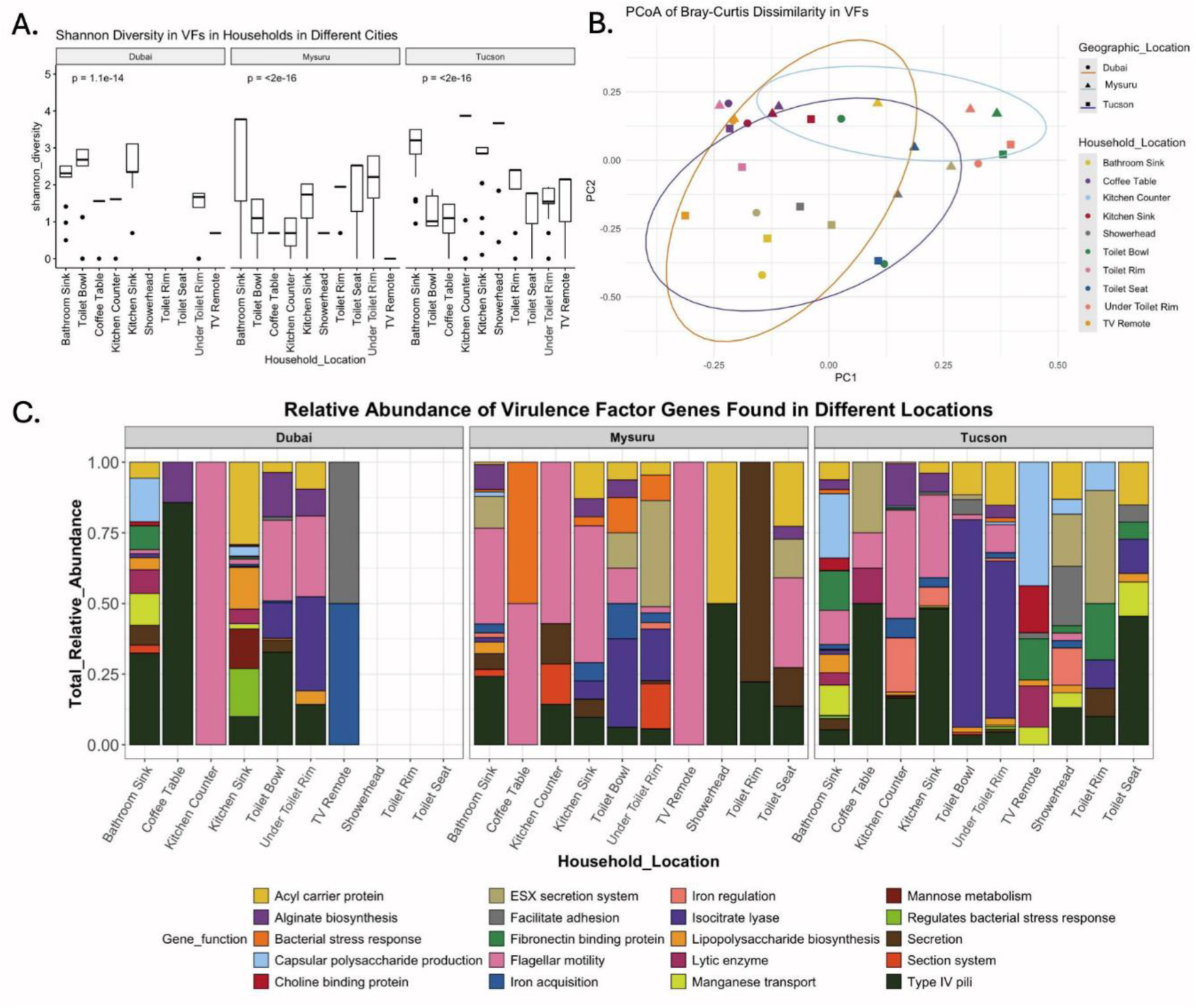
An overview of the virulence factor genes (VFs) found in the ten household locations across three cities. A: Shannon diversity of VFs in household locations, highly significant in all three cities (ANOVA, p<0.01). B: Principal coordinates analysis of Bray-Curtis distances (beta diversity) of ARGs in composite household location samples in the study. Samples are colored by location within the house, point shapes correspond to the geographic location, and ellipses are drawn around clusters by geographic location. PERMANOVA analysis of the samples showed that geographic location and household location were not significant drivers of VF diversity (p=0.129 and p=0.127, respectively). C: The relative abundance of the top 20 most-abundant VF gene functions within the composite household samples in the three cities. No VFs were detected in the Dubai showerhead, toilet rim, or toilet seat composite samples.

**Table 2.** Core virulence genes in each location by household sampling site*.

| City | Kitchen Counter | Kitchen Sink | Bathroom Sink | Overall |
| --- | --- | --- | --- | --- |
| Dubai | None | None | None | None |
| Mysuru | None | None | None | None |
| Tucson | None | None | None | None |
| All | Flagellar motility | Alginate biosynthesis;<br>Flagellar motility;<br>Type IV pili;<br>Iron acquisition;<br>Acyl carrier protein | Isocitrate lyase;<br>Lipopolysaccharide biosynthesis;<br>Capsular polysaccharide production;<br>Flagellar motility;<br>Type IV pili;<br>Secretion;<br>Catalase;<br>Acyl carrier protein;<br>Lipoteichoic acid biosynthesis | Type IV pili;<br>Secretion;<br>Isocitrate lyase;<br>Flagellar motility;<br>Acyl carrier protein |

\* Note: Coffee table, TV remote, toilet seat, toilet rim, toilet bowl, and under the toilet rim did not have any core virulence genes in any location and therefore are not listed on the table.

#### Pathogens

Pathogens were detected across the households and cities, as detailed in **Table 1**. The most abundant pathogens overall were *Escherichia coli*, *Actinobacter baumanii, Klebsiella pneumoniae,* and *Enterobacter* spp. Only one eukaryotic pathogen was detected, in Tucson. The household locations with the greatest diversity of pathogens were the kitchen and bathroom sink, with DNA from 13 to14 unique pathogenic species found in the kitchen sink in each country and DNA from 7 to 10 species on the bathroom sinks. Tucson and Mysuru showed a greater number of unique pathogens across the household overall, but Dubai had a larger number of species under the toilet rim (10 species) and in the kitchen sink (14 species). Viral genera containing pathogens were detected in the samples, but this analysis did not go down to the species level.

#### DNA Virome

Finally, we also examined the virome of the different household locations in the three cities to assess potential DNA viruses present throughout the homes using composite samples from each of the household locations. Variuation was seen throughout the household and between the countries. The DNA virome in Tucson and Dubai were more similar to each other than that of Mysuru. At the viral phylum level, many samples were dominated by Uroviricota, a phylum of double-stranded DNA phages which infect bacteria and archaea, particularly in Mysuru and Dubai. Lenarviricota, a single-stranded RNA phylum which contains both bacteriophages and plant- and fungal-associated viruses(45), was detected in Dubai and Tucson on the kitchen counter and sink, as well as on the bathroom sink and under the toilet rim in Dubai (**Figure 6A**). Dubai and Tucson had more diverse viromes at all taxonomic levels than Mysuru, which was dominated by the bacteriophage Muvirus (**Figure 6E**).

#### Mysuru, India

Mysuru had high levels of aminoglycoside resistance on the kitchen counters and kitchen sinks, macrolide resistance on the toilet bowls, and macrolide, lincosamide, streptogramins type B (MLS-B) resistance on the bathroom sinks and showerheads **(Figure 1C)**. Beta diversity of the ARGs in the Mysuru households could not be explained by differences in within-house locations and was likely influenced by each individual household as a whole (variation was more correlated to house number, ie. house 1, 2, 3… 10) rather than the specific location within the household. Nested PERMANOVA analysis showed a trend very close to significance but was not past the threshold of p=0.05 (p=0.0508). Through a principle coordinates analysis (PCoA), some clustering of ARG profiles can be seen in the kitchen sink, toilet seat, and kitchen counter, however there are no stand-out clusters (**Figure 3A**). The only core ARG between the ten household locations in Mysuru was aminoglycoside resistance (when excluding the TV remotes), suggesting a variable ARG profile was variable between houses and within household locations.

**Figure 3.**
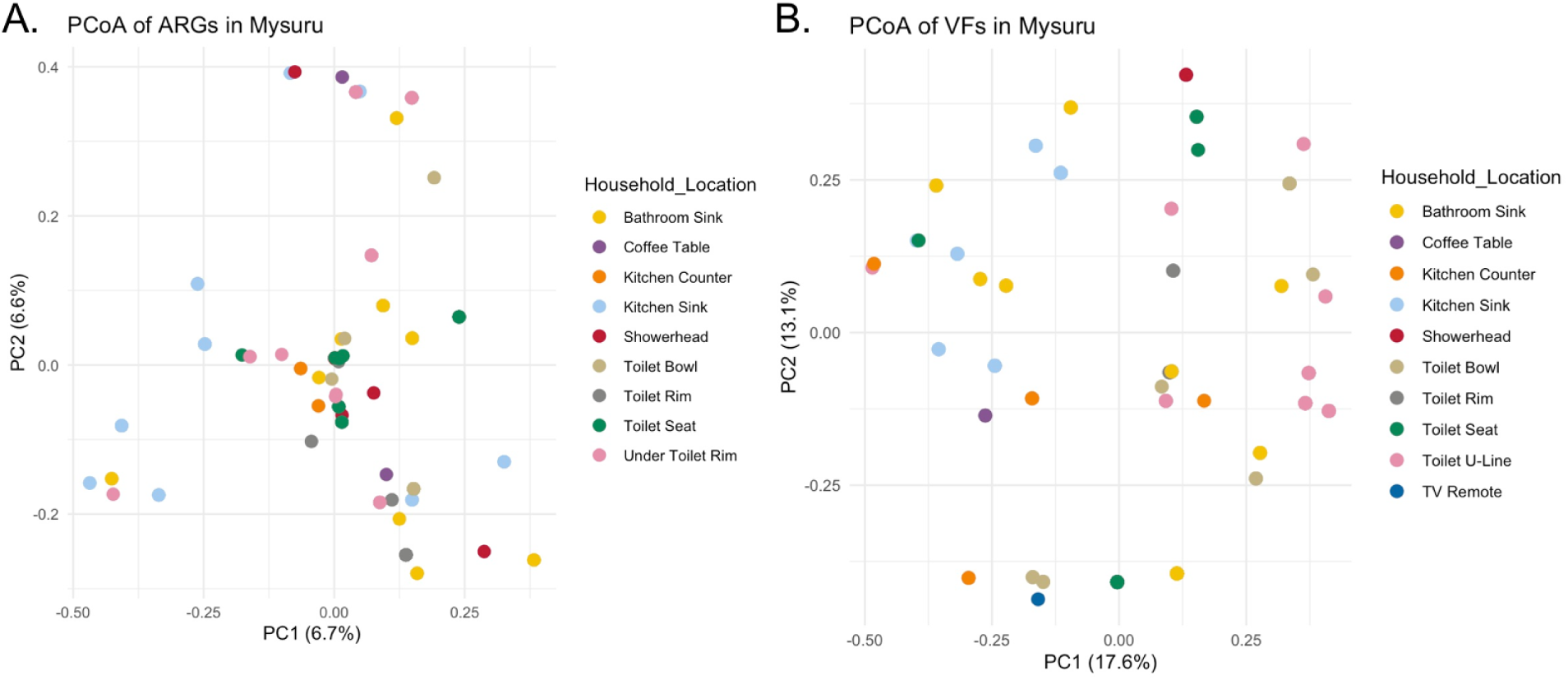
ARG and VF profiles of samples from Mysuru, India. A: PCoA of ARG diversity between the ten household locations from the ten houses in in Mysuru. Within houses, ARGs are not different across the household locations, and differences in the ARG profiles are due to house-to-house variation, not the different locations in the house, although this relationship is approaching significance (PERMANOVA, p=0.0508). B: PCoA of VF diversity between the ten household locations from the ten houses in in Mysuru. VFs are not different across the household locations, and differences in the VF profiles is due to house-to-house variation (PERMANOVA, p=0.1167).

In Mysuru the highest VF diversity was the bathroom sink, toilet seat, under the toilet rim, and kitchen sink (**Figure 2A**). In Mysuru households, flagella-associated genes were highly abundant in the bathroom sink, coffee table, kitchen counter, kitchen sink, TV remote, and toilet seat, while the showerhead had high abundance of both the acyl carrier protein genes and type IV pili (**Figure 2C**). We found that variation in the VF profile was not well explained by the household location or the house number (PERMANOVA, p=0.1167). Our PCoA analysis of the VFs showed minimal clustering of household locations, although more variation in the samples was explained by PC1 and PC2 than in the ARG analysis (**Figure 3B**). Pathogens were detected in all ten household locations in Mysuru. The most commonly detected pathogens in Mysuru were *Acinetobacter baumanii*, found in 48% of samples (taken from the coffee table, kitchen counter and sink, shower head, bathroom sink, and from all four toilet locations), and *Klebsiella pneumoniae* and *Escherichia coli* in 26% of samples (both detected on coffee table, kitchen sink, bathroom sink, showerhead, and all four toilet samples, with *K. pneumoniae* detected on the TV remote and kitchen counter as well). Pathogens were most often identified in the kitchen (34%) and bathroom sinks (18%), and least often found on the TV remote (0.7%), coffee table, and kitchen counter (both contributed to 5%). No eukaryotic pathogens were detected in Mysuru (**Table 3**).

**Table 3.** Potential pathogens in each location by household sampling site.

| Location | Dubai | Mysuru | Tucson |
| --- | --- | --- | --- |
| Coffee Table | *Acinetobacter baumannii* | *Salmonella enterica*;<br>*Escherichia coli*;<br>*Enterobacter cloacae*;<br>*Klebsiella pneumoniae*;<br>*Acinetobacter baumannii*;<br>*Acinetobacter calcoaceticus* | *Escherichia coli*;<br>*Acinetobacter baumannii* |
| TV Remote | *Acinetobacter baumannii* | *Klebsiella pneumoniae* | *Enterobacter hormaechei* |
| Kitchen Counter | *Escherichia coli*;<br>*Enterobacter cloacae*;<br>*Klebsiella pneumoniae*;<br>*Acinetobacter baumannii*;<br>*Enterobacter mori* | *Klebsiella pneumoniae*;<br>*Acinetobacter baumannii* | *Salmonella enterica*;<br>*Escherichia coli*;<br>*Enterobacter hormaechei*;<br>*Enterobacter cloacae*;<br>*Klebsiella pneumoniae*;<br>*Klebsiella oxytoca*;<br>*Enterobacter asburiae*;<br>*Acinetobacter baumannii*;<br>*Enterobacter kobei*;<br>*Enterobacter ludwigii*;<br>*Enterobacter bugandensis*;<br>*Enterobacter mori*;<br>*Enterobacter chuandaensis*;<br>*Enterobacter soli*;<br>*Acinetobacter calcoaceticus* |
| Kitchen Sink | *Salmonella enterica*;<br>*Escherichia coli*;<br>*Enterobacter hormaechei*;<br>*Enterobacter cloacae*;<br>*Klebsiella pneumoniae*;<br>*Klebsiella oxytoca*;<br>*Enterobacter asburiae*;<br>*Acinetobacter baumannii*;<br>*Enterobacter kobei*;<br>*Enterobacter ludwigii*;<br>*Enterobacter bugandensis*;<br>*Enterobacter mori*;<br>*Acinetobacter calcoaceticus*;<br>*Listeria monocytogenes* | *Salmonella enterica*;<br>*Escherichia coli*;<br>*Enterobacter hormaechei*;<br>*Enterobacter cloacae*;<br>*Klebsiella pneumoniae*;<br>*Klebsiella oxytoca*;<br>*Enterobacter asburiae*;<br>*Acinetobacter baumannii*;<br>*Enterobacter kobei*;<br>*Enterobacter ludwigii*;<br>*Enterobacter bugandensis*;<br>*Acinetobacter calcoaceticus*;<br>*Listeria monocytogenes* | *Salmonella enterica*;<br>*Escherichia coli*;<br>*Enterobacter cloacae*;<br>*Klebsiella pneumoniae*;<br>*Klebsiella oxytoca*;<br>*Enterobacter asburiae*;<br>*Acinetobacter baumannii*;<br>*Enterobacter kobei*;<br>*Enterobacter ludwigii*;<br>*Enterobacter bugandensis*;<br>*Enterobacter mori*;<br>*Enterobacter chuandaensis*;<br>*Acinetobacter calcoaceticus* |
| Showerhead | None | *Escherichia coli*;<br>*Klebsiella pneumoniae*;<br>*Acinetobacter baumannii*;<br>*Listeria monocytogenes* | *Escherichia coli*;<br>*Enterobacter hormaechei*;<br>*Enterobacter cloacae*;<br>*Klebsiella pneumoniae*;<br>*Klebsiella oxytoca*;<br>*Acinetobacter baumannii*;<br>*Acinetobacter calcoaceticus* |
| Bathroom Sink | *Escherichia coli*;<br>*Enterobacter hormaechei*;<br>*Enterobacter cloacae*;<br>*Klebsiella pneumoniae*;<br>*Klebsiella oxytoca*;<br>*Acinetobacter baumannii*;<br>*Enterobacter kobei* | *Escherichia coli*;<br>*Enterobacter hormaechei*;<br>*Enterobacter cloacae*;<br>*Klebsiella pneumoniae*;<br>*Acinetobacter baumannii*;<br>*Enterobacter kobei*;<br>*Listeria monocytogenes* | *Salmonella enterica*;<br>*Escherichia coli*;<br>*Enterobacter cloacae*;<br>*Klebsiella pneumoniae*;<br>*Klebsiella oxytoca*;<br>*Acinetobacter baumannii*;<br>*Enterobacter kobei*;<br>*Enterobacter mori*;<br>*Acinetobacter calcoaceticus*;<br>*Coccidioides immitus* |
| Toilet Seat | *Acinetobacter baumannii* | *Escherichia coli*;<br>*Enterobacter hormaechei*;<br>*Enterobacter cloacae*;<br>*Klebsiella pneumoniae*;<br>*Acinetobacter baumannii*;<br>*Acinetobacter calcoaceticus* | *Escherichia coli*;<br>*Klebsiella pneumoniae*;<br>*Klebsiella oxytoca*;<br>*Enterobacter asburiae*;<br>*Acinetobacter baumannii*;<br>*Acinetobacter calcoaceticus* |
| Toilet Rim | None | *Escherichia coli*;<br>*Enterobacter hormaechei*;<br>*Enterobacter cloacae*;<br>*Klebsiella pneumoniae*;<br>*Acinetobacter baumannii* | *Escherichia coli*;<br>*Klebsiella pneumoniae*;<br>*Klebsiella oxytoca* |
| Toilet Bowl | None | *Escherichia coli*;<br>*Klebsiella pneumoniae*;<br>*Acinetobacter baumannii*;<br>*Acinetobacter calcoaceticus*;<br>*Listeria monocytogenes*;<br>*Campylobacter jejuni* | *Salmonella enterica*;<br>*Escherichia coli*;<br>*Enterobacter hormaechei*;<br>*Klebsiella pneumoniae*;<br>*Klebsiella oxytoca*;<br>*Acinetobacter baumannii*;<br>*Acinetobacter calcoaceticus*;<br>*Coccidioides immitus* |
| Under the Toilet Rim | *Escherichia coli*;<br>*Enterobacter hormaechei*;<br>*Enterobacter cloacae*;<br>*Klebsiella pneumoniae*;<br>*Enterobacter asburiae*;<br>*Acinetobacter baumannii*;<br>*Enterobacter kobei*;<br>*Enterobacter bugandensis*;<br>*Acinetobacter calcoaceticus*;<br>*Listeria monocytogenes* | *Escherichia coli*;<br>*Klebsiella pneumoniae*;<br>*Acinetobacter baumannii* | *Salmonella enterica*;<br>*Escherichia coli*;<br>*Enterobacter hormaechei*;<br>*Enterobacter cloacae*;<br>*Klebsiella pneumoniae*;<br>*Klebsiella oxytoca*;<br>*Acinetobacter baumannii*;<br>*Enterobacter kobei*;<br>*Acinetobacter calcoaceticus* |

The DNA virome of Mysuru differed on taxonomic composition from the other cities and was less diverse. At the viral class level, we see Mysuru household locations are dominated by Caudoviricetes, a class of phages in the Uroviricota phylum, which was found at high abundances in Dubai and Tucson as well (**Figure 6B**). The relative viral order abundance demonstrated that all Mysuru locations were dominated by unclassified orders of viruses (**Figure 6C**). At the genus level, Mysuru was greatly dominated by *Muvirus,* a bacteriophage of the class Caudoviricetes (**Figure 6E**).

#### Dubai, UAE

Household samples from Dubai a had high abundance of aminoglycoside resistance in the kitchen sinks, toilet bowls, and under the toilet rims, and macrolide resistance was more abundant in the bathroom sinks, coffee tables, and TV remotes **(Figure 1C)**. The ARG diversity in Dubai households was highly explained by household location, rather than by house-to-house variation (PERMANOVA, p<0.001). The majority of samples cluster together on the PCoA plot, with the samples from under the toilet rim more separate from the rest, indicating a differing ARG profile, and the PC1 and PC2 axes explain a modest amount of variation in the samples (**Figure 4A**).

**Figure 4.**
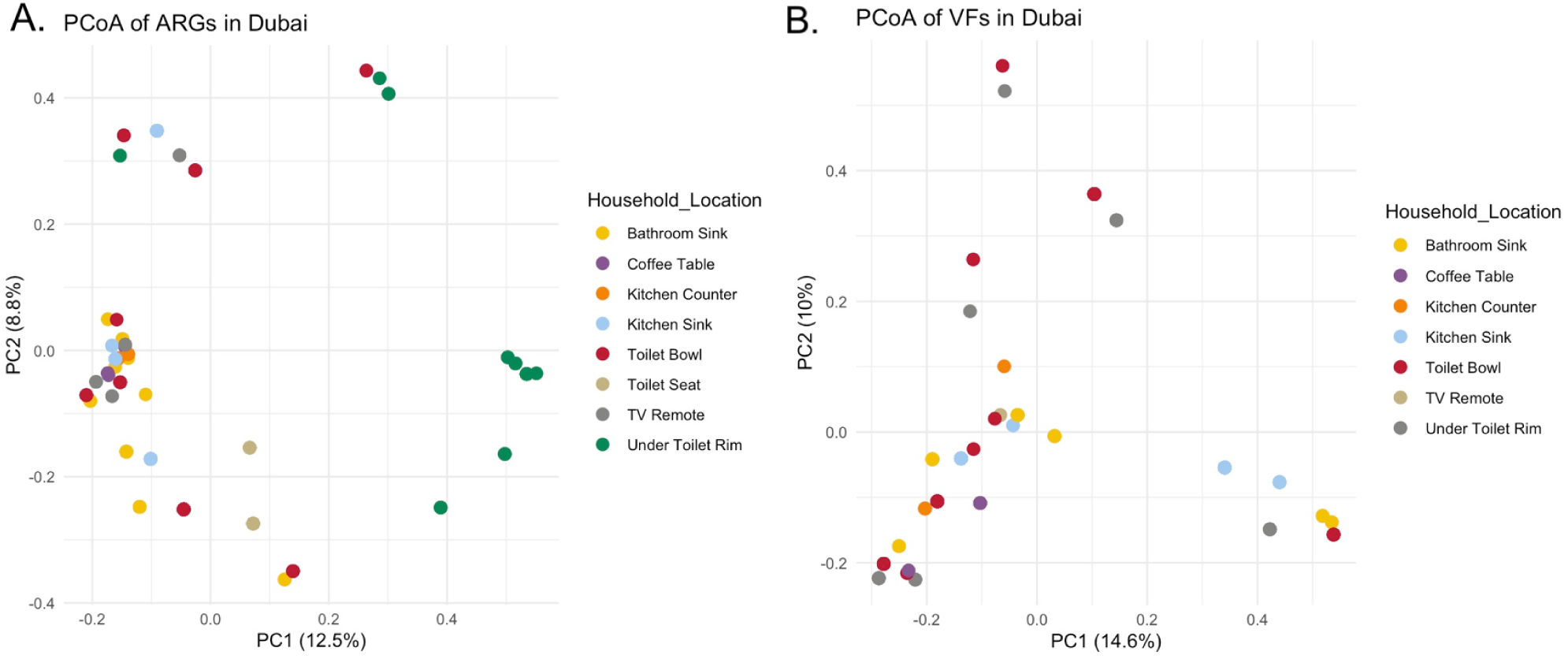
ARG and VF profiles of samples from Dubai, UAE. A: PCoA of ARG diversity between the ten household locations in Dubai using samples from the ten locations in ten houses. Within houses in Dubai, ARGs are very different between household locations regardless of what house the sample came from (PERMANOVA, p<0.001). B: PCoA of VF diversity between seven household locations across the Dubai houses (three locations did not have any detected VFs across all houses in Dubai). Within houses in Dubai, VF diversity is not explained by different locations within the house (PERMANOVA, p=0.0551), but this relationship is approaching significance.

The highest VF diversity in Dubai households was in the toilet bowl, kitchen sink, bathroom sink, and under the toilet rim (**Figure 2A**). The bathroom sink, toilet bowl, and coffee table had a high abundance of type IV pili, and the kitchen counter, toilet bowl, and under the toilet rim had high abundance of flagellar protein genes (**Figure 2C**). The VF profiles were not significantly explained by household location, but were approaching significance (PERMANOVA, p=0.0551). Not all household locations had VFs detected in the samples, and samples did not show clear clustering on a PCoA plot (**Figure 4B**). In Dubai, pathogens were detected in 8 of the ten household locations and were most commonly found on the kitchen sink (32%) and bathroom sink (28%). No pathogens were detected on the toilet seat or rim. Similarly to Mysuru, the most commonly detected pathogens were *Acinetobacter baumanii,* found in 32% of samples (which came from coffee table, TV remote, kitchen counter, kitchen sink, bathroom sink, toilet seat, and under the toilet rim samples), *Klebsiella pneumoniae*, found in 20% of samples (found on kitchen counter, kitchen sink, bathroom sink, and under the toilet rim samples), and *Escherichia coli,* found in 16% of samples which were taken from the kitchen counter and sink, the bathroom sink, and under the toilet rim. Again, no eukaryotic pathogens were detected in the Dubai households (**Table 3**).

At the viral class level, Dubai household locations were also mostly unclassified except for the coffee tables and TV remotes that had high abundance of Chitovirales and the toilet seat which had a higher relative abundance of Crassvirales. Timlovirales was also found in Dubai households on the kitchen counter, bathroom sink, kitchen sink, and toilet bowl and under the toilet rim (**Figure 6C**). The virome of TV remotes and coffee tables were predominately made of the viral family Poxviridae. The toilet seats were dominated by Poxviridae and Intestiviridae, and the kitchen counter and sink, and the bathroom sink and toilet bowl had high levels of Steitzviridae and Poxviridae. Additionally, the toilet rims were dominated by approximately 50% unclassified, ∼30% Poxviridae, and 25% other viruses (**Figure 6D**). Many of the locations in Dubai were mostly classified as ‘other’, but on the bathroom sink, toilet bowl, toilet seat, and showerhead, *Oryzopoxvirus* was more prevalent (**Figure 6E**).

#### Tucson, Arizona, USA

The ARG diversity in Tucson households was significantly influenced by household location when controlling for house-to-house variation (PERMANOVA, p<0.001). PCoA analysis shows clustering of some samples, such as those under the toilet rim and bathroom sink, as well as the coffee table, indicating some consistency in ARG profiles across the ten houses when looking at the same household location, which agrees with the PERMANOVA analysis (**Figure 5A**). In Tucson, beta-lactamases, lincosamide resistance, tetracycline resistance, macrolide resistance, and MLS-B resistance were found at least once in all household locations. The ARG distribution was more variable than the other, but the bathroom sinks had high abundance of tetracycline resistance, on kitchen counters and sinks beta-lactamases were abundant, aminoglycoside resistance was found in the toilet bowls, and MLS-B resistance on the toilet rims and seats (**Figure 1C**).

**Figure 5.**
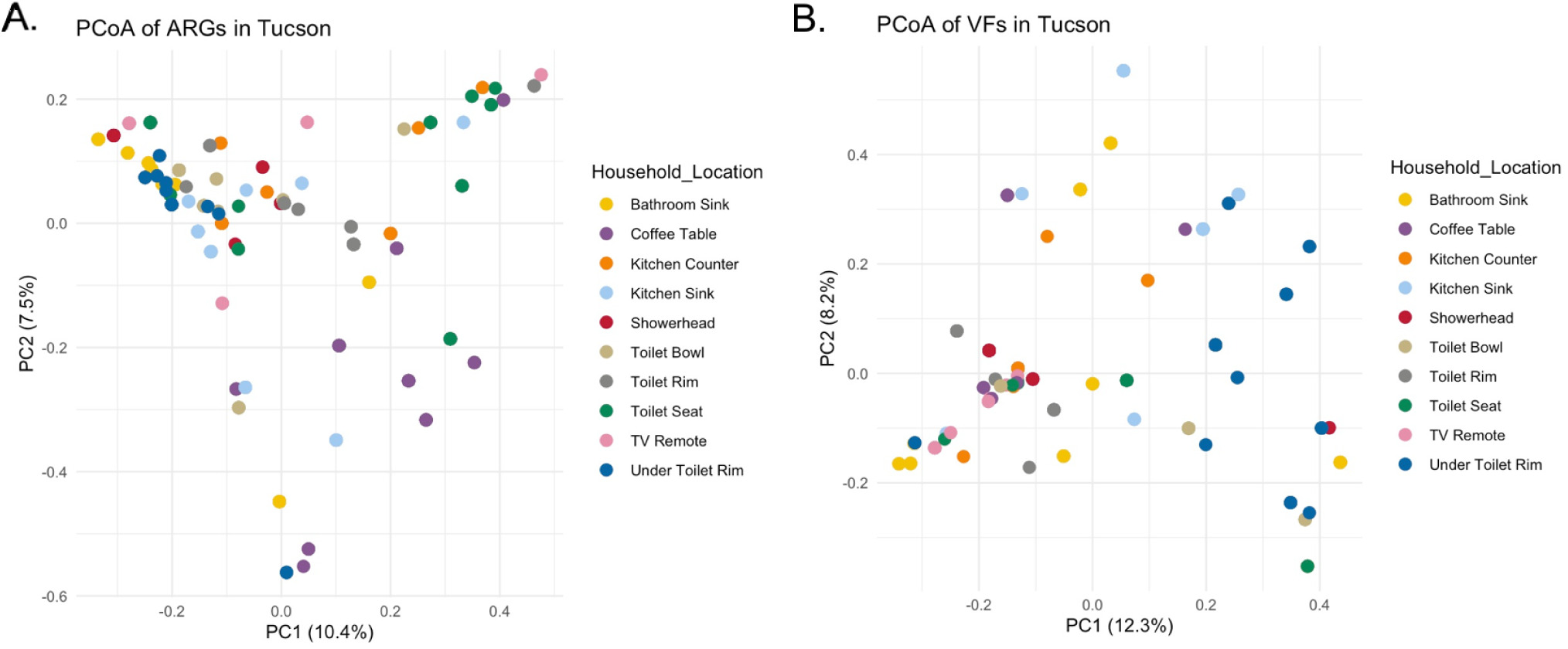
ARG and VF profiles of samples from Tucson, Arizona, USA. A: PCoA of ARG diversity between the ten household locations in Tucson across the ten houses. Within houses in Tucson, ARGs are very different between household locations, and ARG diversity is driven more by location within the house than house-to-house differences (PERMANOVA, p<0.001). B: PCoA of VF diversity between the ten household locations in Tucson. Similarly to ARGs, VF diversity is driven significantly more by household location than house-to-house variation (PERMANOVA, p<0.01).

The highest VF diversity was found on the kitchen counter, bathroom sink, and showerhead (**Figure 2A**). Households in Tucson had a high abundance of isocitrate lyase genes under the toilet rim and in the toilet bowl, and type IV pili on the toilet seats, kitchen sinks, and coffee tables (**Figure 2C**). The VF diversity was also associated with household location (PERMANOVA, p=0.0013), and PCoA analysis showed clustering between the samples from under the toilet rim, the TV remote, and the toilet rim. Similarly to the other countries, the PC1 and PC2 axis explained a small amount of variation in the samples for both the ARGs and VFs (**Figure 5B**). Pathogens were found in all ten household locations in Tucson – 29% of the detected pathogens were in the kitchen sink, 20% in the bathroom sink, and 14% on the kitchen counter. This aligns with pathogen detection in Mysuru and Dubai. The locations with the least pathogens were the TV remote (0.99%), toilet rim (2.4%), and coffee table (1.9%). As with the other two cities, the most common pathogens were *Acinetobacter baumannii* (detected in 37% of samples, coming from the coffee table, kitchen counter and sink, showerhead and bathroom sink, the toilet bowl, and under the toilet rim), *Klebsiella pneumoniae* (detected in 29% of samples, in all household locations except the coffee table and TV remote), and *Escherichia coli* (detected in 36% of samples, again in all locations except the TV remote). In Tucson, one eukaryotic pathogen was found – *Coccidioides immitis*, an endemic soil fungus associated with respiratory disease in the area – which was found 2 of the 100 samples (**Table 3**).

In the Tucson household DNA virome, Nucleocytoviricota was abundant throughout the house, particularly on the coffee table, TV remote, and toilet bowl (it was dominant on the coffee table and TV remote in Dubai as well). These viruses have large DNA genomes, and some members are pathogenic to humans, such as the *Molluscum contagiosum*, which is a skin infection causing raised lesions(46, 47) (**Figure 6A).** Pokkesviricetes was found in high abundances in many Tucson samples, as well as the coffee table and TV remote in Dubai. Leviviricetes and Herviviricetes were found at relative abundances of 25% or less across the samples as well, as the third and fourth most abundant class (**Figure 6B**). Only the bathroom sinks, kitchen sinks, and toilet rims with high abundance of unclassified viral orders in Tucson, while the remaining other household locations were dominated by Chitovirales (**Figure 6C**). At the family level Tucson household locations were dominated by Poxviridae with only the kitchen sinks containing approximately 50% unclassified families and the other 50% Steitzviridae, Poxviridae, Kyanoviridae, and other viruses (**Figure 6D**). In Tucson, *Oryzopoxvirus* was dominant in almost all samples. Low levels of other genera, such as *Kinglevirus* and *Cytomegalovirus* (which is known to be pathogenic to humans), were detected in Tucson (**Figure 6E**).

**Figure 6.**
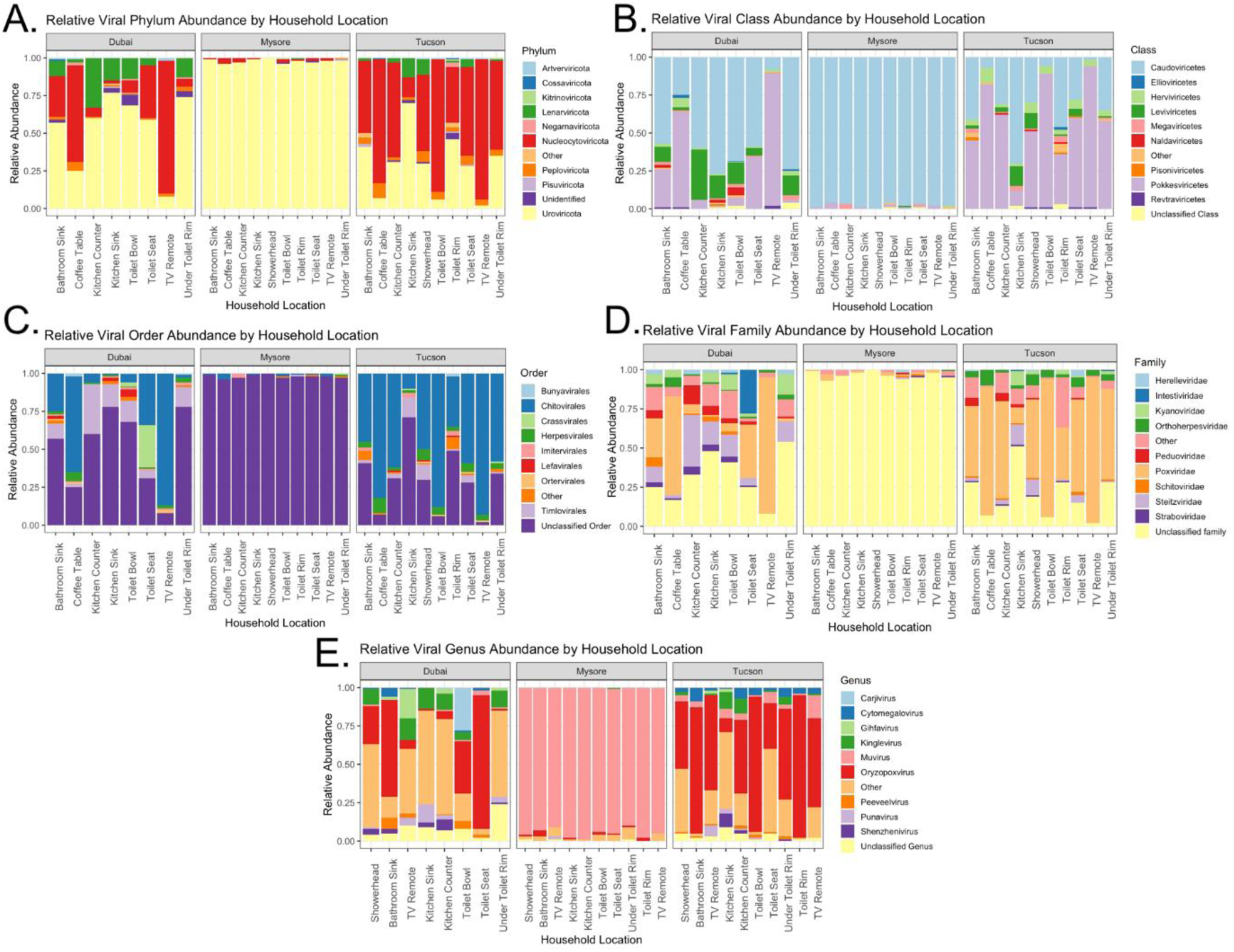
Viral relative abundance at different taxonomic levels. A: Relative abundance of the top ten most abundant phyla in the composite household location samples. B: Relative abundance of the top ten most abundant classes in the composite household location samples. C: Relative abundance of the top ten most abundant order in the composite household location samples. D: Relative abundance of the top ten most abundant families in the composite household location samples. E: Relative abundance of the top ten most abundant genera in the composite household location samples.

## Discussion

This study assessed the prevalence and diversity of ARGs and VFs, as well as the presence of pathogens across three different global locations. There was a high degree of diversity in ARGs and VFs in nearly all samples, with considerable differences between the three cities. Pathogenic bacteria were detected in global locations, most often found in the kitchen and bathroom sinks. The diversity within the genes of the microbiome, particularly between the three cities, emphasizes the role that environmental, cultural, and differences in the socioeconomic status of the country and household occupants may play in shaping the microbiome. This may aid in understanding the composition of the microbiome, why certain genes or taxa favor specific niches, and assessing the risk that the built microbiome poses to human health across the globe. The presence of ARGs within the environment is inevitable(48), but the variability within the categories of resistance, mechanisms, and locations where they are more commonly located can provide information on their role in the built environment. This study found significant differences in the alpha diversity in ARGs across the household in all three cities. Despite the significant differences in the resistome when looking at each household location (across the ten houses in each city) though the Shannon diversity index, when looking at a specific location globally (ie. kitchen counters, no matter whether they are in Mysuru, Tucson, and Dubai), beta diversity analysis revealed that geographic location significantly explained ARG diversity and was a greater driver of the resistome than within-household location. This opposes what was found in other studies, which found a wide variety of ARGs in the built microbiome but the composition of the resistome was more dependent on surface type and usage(14–16). Additionally, Yin et al (2023) looked at the resistome in different environments around the globe (i.e. wastewater from multiple different countries) and found that location type and usage are greater drivers of ARG diversity than geographic location(49). However, these studies only looked at one or two geographic locations (and if looking at multiple locations, they were often relatively close, such as two cities on the eastern side of the United States or countries bordering each other in Europe), so our more diverse geographic locations, spanning across the United States, the Middle East, and Southern Asia may have represented the influence of geography more accurately. Within each city, we found that household location (coffee table, TV remote, etc) explained more of the ARG variation than household number (with highly significant trends in Tucson and Dubai, and a trend approaching significance in Mysuru), suggesting that household location and surface use/type does play an important role in shaping the resistome on a local level. Other studies have found clinically important resistance genes in the built environment, such as resistance to aminoglycosides, MLS-B, polymyxin, bacitracin, and vancomycin(11, 12). Our study found similar results – the most common ARGs across the three countries were against aminoglycosides, MLS-B, and macrolides, however we also saw resistance against tetracycline, particularly in Tucson and Dubai. The core ARG analysis showed a large core resistome across all samples, suggesting that there is potential for the core ARGs to be present across different household locations regardless of geographic location. The core resistome in the built microbiome overall has not been well studied, but these results indicate that it should be further explored to see if any patterns emerge regarding certain resistance genes which may have a more specialized niche in the household, or if the core resistome (ARGs found in the same place within the household across houses and/or cities) is more heavily influenced by geographic location. Antibiotic resistance is a growing issue and the presence of ARGs in the environment suggests that bacteria can potentially pick up these genes and cause infections in humans(2, 8–10), particularly in light of several pathogens detected throughout households in this study. This highlights the need to assess the resistome in more depth to determine the risk and role of ARGs in the built environment. Additionally, knowing that geographic location is a driver of ARG diversity, exploring antibiotic usage in the environment could provide an interesting assessment of how population-wide antibiotic usage (in healthcare, cleaning, and agriculture) and other cultural and environmental factors influence the resistome. This could help determine whether geographic location on a global scale, as we found, plays a stronger role in driving the resistome, or if, as seen in other studies(14–16, 49), surface type or use are the main drivers of ARG diversity in a given built location.

We also explored the presence and abundance of virulence factors, which has not been studied as extensively as the microbiome and ARGs. VFs showed significant levels of variation within each household location in each city, and the beta diversity of VF profiles did not significantly differ between geographic locations or household locations. Both location scales do not appear to be primarily driving VF diversity, so further aspects of the built environment (e.g., cleaning habits, number of occupants) could be explored to identify what is influencing the VF composition. Several studies have identified VFs across the household, typically related to motility, adhesion, and metabolism(11) – this study found similar results, with flagellar motility, type IV pili, and acyl carrier genes being the most common VFs across all samples, along with several other metabolism-related VFs. Locations with the highest diversity of VFs in the three cities were associated with water, such as the kitchen and bathroom sink, and inside the toilet. The presence of moisture in these environments likely leads to the development of biofilms, explaining the higher levels of VF genes, particularly those related to adhesion and motility(50). There were even less core virulence factor genes (VF genes found in the same household and/or geographic location across the set of samples) than ARGs, but a core set of VF genes which were found at least once in any of the household locations in each country was still found, again containing genes related to adhesion, motility, and metabolism. This suggests that an increased ability to move and adhere to household surfaces, and a wider range of metabolic activity may be crucial for survival in the built environment, where nutrient availability may be depleted due to surface characteristics and cleaning habits(4, 23, 51). These associations of certain genes may indicate a niche where these genes are more important to the bacteria carrying them. For example, ARGs and virulence genes for adhesion may be more important for moist environments like sinks, so the bacteria can adhere to the surface despite a frequent influx of water, soaps, and disinfectants that often contain antimicrobials(52). We found that household location was a significant driver of VF diversity in Tucson, approaching significance in Dubai, and not significant in Mysuru, suggesting that VF diverisyt is more nuanced than ARG and is potentially driven by a wider array of factors which were not assessed in this study. The presence of VF genes is important as it indicates the possibility that microorganisms in the environment could be opportunistic pathogens or transfer their genes to increase the virulence of already-present pathogens(8–10). Overall, as with ARGs, the presence of VF genes that are often passed between bacteria indicates the potential for opportunistic pathogens to arise from the built environment, or for the previously detected pathogens to become more virulent, which is a significant public health concern. However, more work should be done to assess the factors driving VF diversity across the built environment both within households and across geographic distances.

The virome of the built environment is also under characterized compared to the bacterial microbiome. This study measured the DNA virome of the household and found variable diversity between the three cities and within the household locations. The Caudoviricites class, which is a class of bacteriophages highly abundant in the human gut(53), was found at very high levels across all samples. This is consistent with findings from a Hong Kong-based study of the built environment(51), and aligns with the idea that the built microbiome is heavily influenced by the human microbiome, as well as potentially suggesting fecal contamination across household surfaces as this class of viruses in abundant in the human GI tract(3). At the genus level, almost all of the DNA virome from Mysuru was *Muvirus,* which is known to be highly transposable – this phage has the potential to pass ARG and VF genes characterized in this study and disseminate them around the environment(54). Interestingly, *Cytomegalovirus* was found to be a part of the top ten most abundant genera in this study, albeit at low levels across the three cities. Stowell et al (2011) found that *Cytolomegalovirus* can survive on surfaces for up to 6 hours on rubber or cloth surfaces and has greater survival on wet surfaces than dry(55). Additionally, Cytomegalovirus persists in the human genome, which may allow for persistent viral shedding into the household environment(56). This virus was found on the bathroom sink, TV remote, and toilet seat composite samples in Dubai and across all ten locations in Tucson, with the highest abundances on the bathroom sink, kitchen counter, and under the toilet rim. This pathogenic virus can cause hearing loss and intellectual disability in infants and is a cause of concern(55). More work should be done to assess the entire virome (RNA and DNA) in order to capture a complete representation of the viruses in the household virome, in order to determine the health risk to those who reside within the house in different countries around the world. Viruses have variable survival rates across surfaces, and it is known that factors like UV exposure, pH, and virus type including body fluid/suspending medium can influence the length of time a virus can survive on a surface(57). More work should be done to determine the driving factors of the viral microbiome, and whether surface type and use are bigger drivers than geographic location in the virome composition. Also, RNA viruses should be characterized more extensively across the global built microbiome.

Finally, the presence of bacterial pathogens in the house was assessed. Many pathogens within the Enterobacteriaceae family were identified across the household, suggesting an increased risk for enteric disease for household occupants. While Awasthi et al, who conducted the only other household microbiome study in India to date, found the most abundant pathogen to be *Actinobacter baumanii*, we detected this species in only eight of our ten study sites but detected *Klebsiella pnuemoniae* on all ten sites throughout the house(13). No previous studies have looked at Dubai households, but we found a range of pathogens throughout the house. The most abundant pathogen in Tucson was *Escherichia coli*, which is consistent with other US-based studies – Carstens et al found *E. coli, K. pneumoniae,* and *Staphylococcus aureus* in the kitchen of Texan households(22). Although Flores et al did not resolve their data to the species level, they detected genera containing pathogens (*Escherichia, Campylobacter, Clostridium,* and *Salmonella*) in their study of kitchens in Colorado, USA(58). However, the abundance of these genera was very low, as it was in our study(58). These findings highlight the need to be cautious and hygienic when cooking and cleaning, as the pathogens can persist on surfaces and spread throughout the entire household. However, by taking proper measures like frequency of disinfection and washing hands with soap or alcohol-based hand sanitizers or rubs, the chances of actually becoming infected by these pathogens can be reduced, and more work should be done to identify factors and surface characteristics that allow for the persistence of these pathogens on household surfaces. Several challenges were faced within this study – firstly, in some locations, there was a very low biomass present after sampling and extracting DNA, which lead to low sequencing yield. This occurred consistently on the same household locations, like the coffee table, TV remote, and toilet seat – all dry, non-porous surfaces, which may explain the limited microbial colonization(59) - and was not associated with geographic location. This made it difficult to explore certain samples in-depth but was remediated by sequencing the samples multiple times and merging samples to form composites, which were used for the relative abundance of ARGs/VFs analysis. Occasionally, cultural differences between the houses lead to differences in the specific sample locations, however an appropriate substitute was always found and samples (i.e. in Tucson, one household used a video game controller as a TV remote, so this was sampled rather than the traditional TV remote). All houses sampled had western toilets and indoor kitchens – these features may not be common across all houses globally and may play a significant role in shaping the ARG, VF, virus, and pathogen profile of the built environment. If surface type or location usage truly is an important driver in gene diversity, the results from a bathroom with a squat toilet or kitchen which is outdoors may have significantly different outcomes. We only collected DNA samples from the surfaces – this limits the virome analysis to viruses with a DNA genome and bacteriophages which have incorporated into bacterial genomes, as RNA virus genomes would not be preserved through the DNA-oriented processing and sequencing protocols, thus limiting the scope of the virome we were capable of measuring. Additionally, DNA sequencing can result in amplification bias, where specific genomes or genes are overrepresented in the sample DNA amplification. This was not controlled for and could impact the results by exaggerating or underrepresenting certain genes in the microbiome. Including negative controls (or sequencing blanks) could also help identify any contamination in the sequence data, as this can be common in low biomass samples. Overall, few logistic issues were encountered in this study and were addressed through small modifications to the sampling or processing protocol to ensure a robust dataset for analysis.

Overall, differences in the ARG and VF profiles across the households and between cities were identified. This suggests that different locations in the household harbor distinct microbial interactions, which is associated with geographic location as well. Certain genes may be more common when they help survival in different niche situations around the household or built environment. Core ARGs and VFs were found throughout the dataset, meaning that some genes can be found throughout the house in all three countries, but there were few instances where a specific gene was highly associated with a household location across all three cities, or where a specific gene was highly associated with all samples from one city. The virome was variable, but many of the viral sequences belonged to bacteriophages. Bacterial pathogens were detected in nearly all household locations in each country, suggesting the indoor microbiome may have the potential to cause health issues, particularly if the pathogens can pick up ARGs and VFs from other microbiota members, and that pathogens can persist in the household niche. This suggests that the indoor microbiome is dynamic, and many genes can contribute to the establishment of the microbiome in the same household location. The ARGs, VFs, bacterial pathogens, and DNA virome found in the built environment should be characterized across more geographic locations and a greater number of external factors like specific cleaning habits, exposure to the outdoor environment should be assessed. Understanding how the ARGs and VFs, as well as viruses and pathogens, establish in specific niches in the built environment is important to determine global trends in the microbiome and how different ecological and human factors may drive these conditions, could help to mitigate any potential public health risks in the built environment from pathogenic genes in the microbiome and from pathogens.

## Materials and Methods

### Household Sampling

Three geographic locations were selected for sample collection: – Tucson (Arizona, USA – population est. 555,000(24)), Mysuru (Karnataka, India – population est. 3,000,000(25)) and Dubai (United Arab Emirates – population est. 3,800,000(26)). Local teams associated with the study recruited the households to participate in the study, and samples were collected in January 2024 (Mysuru), February 2024 (Tucson), and September 2024 (Dubai). Between 2-6 adults and 0-3 children resided in each household, with up to six bathrooms and between two to six bedrooms. In addition to occupant information, details on cleaning frequency and cleaning products used in each house, but this was not considered in the presented analysis (only the geographic and household location were analyzed regarding the presented data). Depending on availability, participant willingness to provide access to the house, and household construction, up to ten samples were collected from each household. In total, 100 samples were collected in both Tucson and Mysuru and 89 samples were collected in Dubai. The following locations were sampled: kitchen counter surface, kitchen sink (surface and around but not inside the drain), coffee table surface, TV remote (all sides and buttons), showerhead, bathroom sink (surface and around but not inside the drain), and four locations on the toilet – the seat (outside and inside), rim of the toilet (top and outside), toilet bowl above the water, and under the toilet rim (the area underneath the rim inside the bowl). Dual-end swabs (Becton Dickinson, BBL CultureSwab) were used to collect samples - these were dipped in a sterile detergent solution (0.15 M NaCl with 0.1% Tween 20)(27) and then dragged across the sample surfaces for 15 seconds or more while the swab was rotated to collect microbes across the entire surface area. The wide amount of physical variation between households made it difficult to completely standardize collection – to reduce variation in the same collection the samples were taken from the entire surface of each sample site (i.e., corners, edges, center, etc.). Swabs were kept in their original sterile tubes and stored on ice in a cooler for less than 8 hours prior to storing at -20°C long term.

Before conducting the study, Tucson and Dubai study designs were reviewed by the University of Arizona’s Institutional Review Board (IRB). The IRB determined that no IRB was necessary because no human identifying data was collected. The study design for Mysuru households was approved by the Public Health Research Institute of India’s IRB Committee before sampling began.

### DNA Extraction

One of the two swabs was removed from the storage tube and DNA was extracted using the DNeasy Powersoil Pro Kit (Qiagen) per the manufacturer’s instruction with a single modification – samples were heated at 65°C for 10 minutes after adding the first lysis buffer (CD1)(28). DNA concentration was measured using the Qubit fluorometer with the High-Sensitivity Assay (Invitrogen). If the DNA concentration too low to detect using the Qubit assay, DNA was extracted from the second swab to increase yield.

### Oxford Nanopore Sequencing

Sequencing libraries for Dubai and Tucson samples were prepared using the Oxford Nanopore Rapid PCR Barcoding Kit 24 V14 (Oxford Nanopore Technology, SQK-RPB114.24) per the manufacturer’s instructions in the Cooper laboratory. Sequencing libraries for India were prepared at PHRII using the Oxford Nanopore Rapid Barcoding Kit 96 V14 (Oxford Nanopore Technology, SQK-RBK114.96) according to the manufacturer’s protocols. All samples were sequenced using Oxford Nanopore Technology (ONT) on MinION Mk1B devices (Oxford Nanopore Technology) using R10.4.1 flow cells (Oxford Nanopore Technology). Multiple rounds of sequencing were done on each sample library to increase the data yield from each sample.

### Taxonomic Analysis

Reads were concatenated across sequencing runs to form a single fastq file for each of the 289 sampling locations. Composite reads per each household location in each city were generated by concatenating reads from each location across the ten houses in each city (i.e. all kitchen counters in Tucson). Reads were analyzed for bacterial and viral taxonomic composition using the Kraken2(29) software (v2.3.1). The database used in Kraken2 contained all NCBI reference genomes and all non-redundant sequences; this was created by the Cooper laboratory following the kraken2-build protocol for Kraken2. This analysis was done to assess the virome of the samples and to identify bacterial pathogens of interest. Kraken2 was run a second time using the EuPathDB-28, (which contains cleaned eukaryotic pathogen genomes) to detect eukaryotic pathogens(30). Kraken2 was run on all concatenated fastq files (non-composites and composites) with the basic parameters. A BIOM-format file was made using the kraken_biom(31) software (v1.2.0) from the Kraken2 outputs from the viral database. This was analyzed in R using the package phyloseq (v1.52.0), to determine viral taxonomy(32). Viral reads were rarefied to a sample size of 100, as several reads had low biomass in order to not drop any samples. The bacterial and eukaryotic pathogen Kraken2 outputs were used to assess the presence of target pathogens in the samples using a python script. The viral-filtered phyloseq object was used for the viral taxonomic makeup using the following packages: phyloseq, microbiome (v1.30.0)(33), microViz (v0.12.7)(34), and vegan (v2.7.3)(35).

### Antibiotic Resistance Genes and Virulence Factors

Antibiotic resistance genes were identified using Abricate (1.0.1)(36) with the ResFinder(37) database (last updated November 4^th^ 2023), and virulence factors were assessed using Abricate with VFDB(38) (Virulence Factor Database, last updated November 4^th^ 2023)). Abricate identifies ARGs and VF genes from FASTQ files by referencing a pre-installed database and matching genes with a minimum of 15% gene coverage as DNA was not assembled prior to Abricate analysis. Abricate outputs a tab-deliminated file with the identified genes and coverage, and all files were combined into a master Microsoft Excel spreadsheet using python scripts (see Github link). Reads were standardized in Microsoft Excel by dividing the total number of reads per sample by 1,000, and hits from each database were scaled against the standardized reads (for example, if a sample had 100,000 reads and 15 ARG hits, the scaled number of ARG hits would be (100,000 / 1,000) / 15, or 6.67). Samples with a higher number of reads inherently had a larger number of identified VFs or ARGs, which was accounted for by standardizing the hits per reads. Standardized ARG/VF data was imported into R studio and the relative abundance of the top 20 most-abundant ARGs and VFs were plotted for each composite household location sample in each city. To assess the alpha and beta diversity of ARGs and VFs, the relative abundance of each detected gene was calculated across all non-composite samples using dpylr (v1.2.0) in R. This was done in order to assign a scaled value of abundance for every gene across all samples to look at the distribution of each gene. The relative abundances were used to calculate Shannon diversity and Bray-Curtis dissimilarities using the microbiome package in R. ANOVA tests were used to identify significant differences in Shannon diversity across the non-composite samples (n=289) and PERMANOVA tests were used to identify significant differences in beta diversity based on geographic and household locations in the composite samples (n=28). Bray-Curtis dissimilarity values were calculated for ARGs and VFs found in each city individually, which were analyzed with PERMANOVA tests assessing differences in the household location and between the households within each city. This analysis used the R packages readxl(39) (to import data) (v1.4.5), tidyverse (v2.0.0)(40), broom (v1.0.12)(41), rstatix (v0.7.3)(42), reshape2 (v1.4.5)(43), and dpylr (to edit data structure and add metadata), microbiome was used to run the alpha and beta diversity calculations, and ggplot2 (v4.0.2) and ggpubr (v0.6.3) were used for visualization. A ‘core’ ARG and VF is defined by a set of genes present in all samples from a specific location, whether it be geographic (eg. An ARG found in all samples from Dubai would be a part of the core Dubai resistome, or a VF found on every bathroom sink would be a part of the core virulence factor gene to the bathroom sink). The core ARGs and VFs were determined using an online Venn diagram tool (https://bioinformatics.psb.ugent.be/webtools/Venn/) by VIB and UGent (Belgium). Lists of the genes found in all datasets for each particular analysis in a subset of the data were pasted in, and the center of the Venn diagram represented the core genes in that dataset.

### Pathogen Detection

Pathogen detection was done via a python script (see Github link). To detect pathogens, Kraken2 outputs, created using the entire NCBI taxonomic database including non-redundant sequences, the viral kraken2 database, and the eukaryotic pathogen database were scanned for the NCBI taxonomic IDs of specific pathogens of interest. NCBI taxonomic IDs were found via the NCBI taxonomy browser and were specified to the species level(44). Kraken2 outputs list the number of kmers which match to a specific NCBI taxonomic ID in order of the sequence – outputs were firstly searched for the specified taxonomic IDs (listed in **Supplementary Table 1**). If the pathogen was found in 3 or more kmers (35-mers) in a single read, it was determined to be present in the sample.

### Statistical analysis

All statistical tests were conducted in RStudio. Tests were statistically significant if p was less than or equal to 0.05. Analysis of alpha diversity metrics for the ARG and VF profiles was done using the Shannon index, measuring richness and evenness of genes in each sample. Statistical differences in alpha diversity were calculated using ANOVA tests, to compare between household locations or geographic locations. Beta diversity was analyzed using Bray-Curtis dissimilarity, and, when applicable, PERMANOVA was used to compare Bray-Curtis dissimilarity between geographic locations as the data was non-normally distributed. The formulas used for PERMANOVA calculations were as follows: (dist_matrix ∼ geographic_location) was used to determine differences between the geographic locations alone, and (dist_matrix ∼ household_location) was used to determine differences between household locations alone (for both ARGs and VFs). A nested PERMANOVA was also run on the Bray-Curtis dissimilarity matrix for ARGs and VFs, where household location was nested under geographic location; the formula used was (dist_matrix ∼ household_location + geographic_location) with the “by=”margin”’ argument. For each city individually, nested PERMANOVAs were run on the ARGs and VFs, with household locations (kitchen sink, coffee table, etc) stratified by household (house one, house two) using the formula (dist_matrix ∼ household_location) with the “strata = metadata$house_number” and the “by = “margin”’ argument to account for household differences. This was done to determine whether gene profile differences were driven by differences in the household locations which were consistent across the ten houses, or by differences unique to each individual house. All PERMANOVA tests were run using the adonis2 function from vegan (v2.7.3)(35) in R.

### Data Availability and Visualization

Both Microsoft Excel and RStudio were used to reorganize and subset data, depending on which was more straightforward in each case. Results were represented using a variety of visual plots, generated using ggplot2 in R (version 4.3.5) in RStudio (version 2024.04.0+735). Code used for data analysis can be found on GitHub (https://github.com/carolinescranton01/Global_Household_Microbiome). The sequencing reads from this project can be accessed through the NCBI Bioproject PRJNA1416920.

## Data Availability

DNA sequence data from this research is publicly available on the NCBI’s Sequence Read Archive under BioProject PRJNA1416920. Data was analyzed using python and R code. Analysis protocols and information on software versions, packages, and more can be found within the text and in the following github repository: https://github.com/carolinescranton01/Global_Household_Microbiome. **The authors confirm all supporting data, code and protocols have been provided within the article or through supplementary data files.**

## Acknowledgements

Thank you to the volunteers in Tucson, Dubai, and Mysuru who generously allowed for sample collection from their household.

## References

1. Benton L, Lopez-Galvez N, Herman C, Caporaso G, Cope E, Rosales C, et al. Environmental and Structural Factors Associated with Bacterial Diversity in Household Dust Across the Arizona-Sonora Border. Res Sq. 2023.

2. Jeon YS, Chun J, Kim BS. Identification of household bacterial community and analysis of species shared with human microbiome. Curr Microbiol. 2013;67(5):557–63.

3. Rai S, Singh DK, Kumar A. Microbial, environmental and anthropogenic factors influencing the indoor microbiome of the built environment. J Basic Microbiol. 2021;61(4):267–92.

4. National Academies of Sciences Eg, and Medicine, Engineering NAo, Sciences DoEaP, Division HaM, Studies DoEaL, Environment BoIatC, et al. Microbiomes of the Built Environment: A Research Agenda for Indoor Microbiology, Human Health, and Buildings. 2017.

5. Klepeis NE, Nelson WC, Ott WR, Robinson JP, Tsang AM, Switzer P, et al. The National Human Activity Pattern Survey (NHAPS): a resource for assessing exposure to environmental pollutants. J Expo Anal Environ Epidemiol. 2001;11(3):231–52.

6. Ferguson L, Taylor J, Davies M, Shrubsole C, Symonds P, Dimitroulopoulou S. Exposure to indoor air pollution across socio-economic groups in high-income countries: A scoping review of the literature and a modelling methodology. Environ Int. 2020;143:105748.

7. Dunn RR, Fierer N, Henley JB, Leff JW, Menninger HL. Home life: factors structuring the bacterial diversity found within and between homes. PLoS One. 2013;8(5):e64133.

8. Zhou JC, Wang YF, Zhu D, Zhu YG. Deciphering the distribution of microbial communities and potential pathogens in the household dust. Sci Total Environ. 2023;872:162250.

9. Carrazana E, Ruiz-Gil T, Fujiyoshi S, Tanaka D, Noda J, Maruyama F, et al. Potential airborne human pathogens: A relevant inhabitant in built environments but not considered in indoor air quality standards. Sci Total Environ. 2023;901:165879.

10. Arnold BJ, Huang IT, Hanage WP. Horizontal gene transfer and adaptive evolution in bacteria. Nat Rev Microbiol. 2022;20(4):206–18.

11. Zhou ZZ, Zhu J, Yin Y, Ding LJ. Seasonal variations of profiles of antibiotic resistance genes and virulence factor genes in household dust from Beijing, China revealed by the metagenomics. Sci Total Environ. 2024;928:172542.

12. Ding LJ, Zhou XY, Zhu YG. Microbiome and antibiotic resistome in household dust from Beijing, China. Environ Int. 2020;139:105702.

13. Awasthi S, Hiremath VM, Nain S, Malik S, Srinivasan V, Rose P, et al. Microbial landscape of Indian homes: the microbial diversity, pathogens and antimicrobial resistome in urban residential spaces. Environ Microbiome. 2025;20(1):25.

14. Du S, Lin H, Luo Q, Man CL, Lai SH, Ho KF, et al. House dust microbiome differentiation and phage-mediated antibiotic resistance and virulence dissemination in the presence of endocrine-disrupting chemicals and pharmaceuticals. Microbiome. 2025;13(1):96.

15. Marshall BM, Robleto E, Dumont T, Levy SB. The frequency of antibiotic-resistant bacteria in homes differing in their use of surface antibacterial agents. Curr Microbiol. 2012;65(4):407–15.

16. Mahnert A, Moissl-Eichinger C, Zojer M, Bogumil D, Mizrahi I, Rattei T, et al. Man-made microbial resistances in built environments. Nat Commun. 2019;10(1):968.

17. Liao H, Liu C, Zhou S, Eldridge DJ, Ai C, Wilhelm SW, et al. Prophage-encoded antibiotic resistance genes are enriched in human-impacted environments. Nat Commun. 2024;15(1):8315.

18. Goh S. Phage Transduction. Methods Mol Biol. 2016;1476:177–85.

19. Rosario K, Fierer N, Miller S, Luongo J, Breitbart M. Diversity of DNA and RNA Viruses in Indoor Air As Assessed via Metagenomic Sequencing. Environ Sci Technol. 2018;52(3):1014–27.

20. Du S, Tong X, Lai ACK, Chan CK, Mason CE, Lee PKH. Highly host-linked viromes in the built environment possess habitat-dependent diversity and functions for potential virus-host coevolution. Nat Commun. 2023;14(1):2676.

21. Wißmann JE, Kirchhoff L, Brüggemann Y, Todt D, Steinmann J, Steinmann E. Persistence of Pathogens on Inanimate Surfaces: A Narrative Review. Microorganisms. 2021;9(2).

22. Carstens CK, Salazar JK, Sharma SV, Chan W, Darkoh C. Evaluation of the kitchen microbiome and food safety behaviors of predominantly low-income families. Front Microbiol. 2022;13:987925.

23. Thompson JR, Argyraki A, Bashton M, Bramwell L, Crown M, Hursthourse AS, et al. Bacterial Diversity in House Dust: Characterization of a Core Indoor Microbiome. *Front*. Environ. Sci; 2021.

24. cenus.gov. Tucson city, Arizona. data.cenus.gov: The United States Cenus Bureau.

25. Lakshmikanth Reddy.G IAS. Mysuru District Website. District At a Glance.

26. Dubai Data and Statistics Establishment. Government of Dubai; 2024.

27. Leff JW, Fierer N. Bacterial communities associated with the surfaces of fresh fruits and vegetables. PLoS One. 2013;8(3):e59310.

28. Berg-Lyons D, Lauber C, Humphrey G, Thompson L, Gilbert J, Jansson J, et al. EMP DNA Extraction Protocol. Protocols.io: EMP Consortium; 2018.

29. Wood DE, Lu J, Langmead B. Improved metagenomic analysis with Kraken 2. Genome Biol. 2019;20(1):257.

30. Lu J, Salzberg SL. Removing contaminants from databases of draft genomes. PLoS Comput Biol. 2018;14(6):e1006277.

31. SM D. kraken-biom: Enabling interoperative format conversion for Kraken results (Version 1.2). 2016.

32. McMurdie PJ, Holmes S. phyloseq: an R package for reproducible interactive analysis and graphics of microbiome census data. PLoS One. 2013;8(4):e61217.

33. Lahti L, <https: 0000-0001-5537-637x=”” orcid.org=””><https: 0000-0001-5537-637x=”” orcid.org=””>Shetty S. microbiome: Microbiome Analytics V1.28.0. github2024.

34. Barnett DJM, Arts ICW, Penders J. microViz: an R package for microbiome data visualization and statistics. Journal of Open Source Software: The Open Journal; 2021. p. 3201.

35. J O, G S, F B, R K, P L, P M, et al. vegan: Community Ecology Package. R package version 2.6-6.1 ed2024.

36. T S. Abricate. Github2020.

37. Zankari E, Hasman H, Cosentino S, Vestergaard M, Rasmussen S, Lund O, et al. Identification of acquired antimicrobial resistance genes. J Antimicrob Chemother. 2012;67(11):2640–4.

38. Chen L, Zheng D, Liu B, Yang J, Jin Q. VFDB 2016: hierarchical and refined dataset for big data analysis--10 years on. Nucleic Acids Res. 2016;44(D1):D694–7.

39. H W, J B. readxl: Read Excel Files. R package version 143 2023.

40. Wickham H, Averick M, Bryan J, Chang W, McGowan LDA, François R, et al. Welcome to the {tidyverse}. Journal of Open Source Software2019. p. 1686.

41. Robinson D, Hayes A, Couch S. broom: Convert Statistical Objects into Tidy Tibbles. 2024.

42. Kassambara A. rstatix: Pipe-Friendly Framework for Basic Statistical Tests. 2023.

43. Wickham H. Reshaping Data with the reshape Package. Journal of Statistical Software2007. p. 1–20.

44. Schoch CL, Ciufo S, Domrachev M, Hotton CL, Kannan S, Khovanskaya R, et al. NCBI Taxonomy: a comprehensive update on curation, resources and tools. Database (Oxford). 2020;2020.

45. Sadiq S, Chen YM, Zhang YZ, Holmes EC. Resolving deep evolutionary relationships within the RNA virus phylum. Virus Evol. 2022;8(1):veac055.

46. Chen X, Anstey AV, Bugert JJ. Molluscum contagiosum virus infection. Lancet Infect Dis. 2013;13(10):877–88.

47. Witt ASA, Carvalho JVRP, Serafim MSM, Arias NEC, Rodrigues RAL, Abrahão JS. The GC% landscape of the Nucleocytoviricota. Braz J Microbiol. 2024;55(4):3373–87.

48. Larsson DGJ, Flach CF. Antibiotic resistance in the environment. Nat Rev Microbiol. 2022;20(5):257–69.

49. Yin X, Li L, Chen X, Liu YY, Lam TT, Topp E, et al. Global environmental resistome: Distinction and connectivity across diverse habitats benchmarked by metagenomic analyses. Water Res. 2023;235:119875.

50. Sharma S, Mohler J, Mahajan SD, Schwartz SA, Bruggemann L, Aalinkeel R. Microbial Biofilm: A Review on Formation, Infection, Antibiotic Resistance, Control Measures, and Innovative Treatment. Microorganisms. 2023;11(6).

51. Leung MHY, Tong X, Lee PKH. Indoor Microbiome and Airborne Pathogens. Comprehensive Biotechnology2019. p. 96–106.

52. Halden RU. On the need and speed of regulating triclosan and triclocarban in the United States. Environ Sci Technol. 2014;48(7):3603–11.

53. Gulyaeva A, Garmaeva S, Kurilshikov A, Vich Vila A, Riksen NP, Netea MG, et al. Diversity and Ecology of. Viruses. 2022;14(10).

54. Harshey RM. Transposable Phage Mu. Microbiol Spectr. 2014;2(5).

55. Stowell JD, Forlin-Passoni D, Din E, Radford K, Brown D, White A, et al. Cytomegalovirus survival on common environmental surfaces: opportunities for viral transmission. J Infect Dis. 2012;205(2):211–4.

56. Goodrum F, Caviness K, Zagallo P. Human cytomegalovirus persistence. Cell Microbiol. 2012;14(5):644–55.

57. P V, I P, M V, A C. Issues Concerning Survival of Viruses on Surfaces. Food Environ Virol: Springer Nature; 2010. p. 24–34.

58. Flores GE, Bates ST, Caporaso JG, Lauber CL, Leff JW, Knight R, et al. Diversity, distribution and sources of bacteria in residential kitchens. Environ Microbiol. 2013;15(2):588–96.

59. Kramer A, Schwebke I, Kampf G. How long do nosocomial pathogens persist on inanimate surfaces? A systematic review. BMC Infect Dis. 2006;6:130.

